# Calculation of centralities in protein kinase A

**DOI:** 10.1101/2022.01.03.474877

**Authors:** Alexandr P. Kornev, Phillip C. Aoto, Susan S. Taylor

## Abstract

Topological analysis of amino acid networks is a common method that can help to understand the roles of individual residues. The most popular approach for network construction is to create a connection between residues if they interact. These interactions are usually weighted by absolute values of correlation coefficients or mutual information. Here we argue that connections in such networks have to reflect levels of cohesion within the protein instead of a simple fact of interaction between residues. If this is correct, an indiscriminate combination of correlation and anti-correlation, as well as the all-inclusive nature of the mutual information metrics, should be detrimental for the analysis. To test our hypothesis, we studied amino acid networks of the protein kinase A created by Local Spatial Pattern alignment, a method that can detect conserved patterns formed by C_α_-C_β_ vectors. Our results showed that, in comparison with the traditional methods, this approach is more efficient in detecting functionally important residues. Out of four studied centrality metrics, Closeness centrality was the least efficient measure of residue importance. Eigenvector centrality proved to be ineffective as the spectral gap values of the networks were very low due to the bilobal structure of the kinase. We recommend using joint graphs of Betweenness centrality and Degree centrality to visualize different aspects of amino acid roles.

Author Summary

Protein structures can be viewed as networks of residues with some of them being a part of highly interconnected hubs and some being connectors between the hubs. Analysis of these networks can be helpful for understanding of possible roles of single amino acids. In this paper, we challenged existing methods for the creation of such networks. A traditional way is to connect residues if they can interact. We propose that residues should be connected only if they retain their mutual positions in space during molecular dynamic simulation, that is they move cohesively. We show that this approach improves the efficiency of the analysis indicating that a significant revision of the existing views on amino acid networks is necessary.

## Introduction

Complex networks are ubiquitous and are studied by diverse fields of science -from physics and biology to sociology and cosmology. Complex networks, unlike random networks or regular lattices, have a modular distribution of their elements (1). Such structural heterogeneity implies that different elements of complex networks have different “importance”. That brings up a graph theoretical problem of network centrality, i.e., a metrics that can quantify this “importance”. Obviously, the definition of “important” elements depends on the context. It can be super spreaders in a pandemic (2), vulnerability points of transportation networks (3), or essential proteins in a metabolic pathway map (4). First attempts to analyze connectivity in social networks were made as early as 1950-60s (5, 6) but the concept of centrality was clarified and formalized in a set of influential papers of Freeman and coworkers in the 1980s (7-9). Since then, multiple centrality measures have been introduced and analyzed in different networks; however, there is no consensus on what centralities should be used for different networks (10). For example, it was shown that centrality measures can be correlated or not correlated depending on the nature of the network (10, 11). It is, therefore, important to study networks on a case-by-case basis with proper benchmark data that can serve as a guide for the selection of efficient centrality metrics.

In this paper, a set of centrality measures has been applied to Protein kinase A (PKA) to find the most effective methods to detect residues that are important for protein function and its regulation. The selection of PKA as an object for this study was dictated by three reasons: First – protein kinases represent one of the largest and crucially important protein families (12) that makes kinases an attractive therapeutic targets (13). Understanding the mechanism of their function and regulation is essential for the rational design of protein kinase inhibitors. Second – protein kinases are dynamic molecules (14), whose function can be modified by distant signaling when a small inhibitor or another protein binds far from the active site (15-17). Such long-distance signaling, known as allostery, is a universal feature in molecular biology (18, 19). Understanding regulatory mechanisms in protein kinases can also be beneficial for studies of other allosteric enzymes. Finally, the third reason is that PKA is, arguably, the most studied protein kinase with a large volume of experimental data accumulated in the past thirty years (20, 21). Therefore, it is easy to create a benchmark for this study, as a large set of key residues that play important functional and regulatory roles are well known.

Multiple authors have previously used graph theoretical approach to analyze different protein properties. It was shown that Closeness centrality can detect residues that are critical for folding of Chymotrypsin inhibitor 2 and the SH3 domain of C-Src kinase (22). A study of 178 protein structures, including ERK2 MAP protein kinase, showed that active site residues have higher Closeness centrality but not Degree centrality (23). Another work, confirming a high level of Closeness centrality of active site residues was done using structures of 46 different families of proteins (24). A study of Voltage-gated sodium channel NaV1.7 showed that residues related to gain-of-function mutations are associated with significant changes of their Betweenness centrality, but not their Closeness centrality, Degree centrality, or Eccentricity (25). A computational study of the Hsp90 chaperone found that residues with high Betweenness centrality are implicated in allosteric communication within the molecule (26). Similar results were reported for G-protein coupled receptor A_2A_AR (27). The authors found that Betweenness centrality was significantly more effective in detecting functionally important residues than Degree or Closeness centralities. A study of allostery in Imidazole glycerol phosphate synthase showed that Eigenvector centrality can detect regions responsible for allosteric communication (28).

It should be pointed out that a possible source of contradictions between these results can lie in the different methods used for the network construction. Currently, there is not even an established term for these networks. Known, in particular, as Protein Contacts Networks (29), Amino Acid Networks (30), or Dynamic Residue Networks (31) they use different criteria for creating links between nodes in the graph. Generally, nodes, representing amino acid residues, are connected if the corresponding residues interact with each other. Different criteria can be used to define the interaction condition: a certain cutoff distance between C_α_ atoms (32), C_β_ atoms (33), heavy atoms of the side chain (34), or centers of mass of the side chain (35). The value of the cutoff varies from 5-9Å (36) to 12-15Å (37) depending on the system but usually is restricted to direct contacts between residues. The links can be weighted using different metrics, such as estimated energy of interaction (38) or absolute value of cross-correlation (34, 39, 40). Treating anti-correlation as correlation is usually justified by the argument that the sign of interaction between residues is not important as soon as information is transmitted between them. A similar argument was made when mutual information was introduced as a measure for interaction between residues in proteins (41). Mutual information is currently widely used in protein network studies (28, 42, 43).

Although merging all interactions irrespective of their type or sign seems to be justified, it can be argued that this approach can be counterproductive and should be reevaluated. Originally, social network analysis has been developed to dissect a set of people into separate groups with shared interests, traits or activities (44). Although, initially, the concept of interaction was intuitively obvious and included only positive relations, such as acquaintance or cooperation, it was subsequently recognized that in certain networks negative interactions such as aggression or hate can be essential for the analysis (45-47). In line with the original social network community analysis, we hypothesized that connectivity in proteins should reflect not a fact of interaction due to spatial proximity of amino acids, but whether they share a dynamic feature, namely, they move cohesively and behave as a semi-rigid body. This is not a new idea, as there are multiple protein dynamics studies that consider rigid communities in proteins (48-50). It has, however, led us to the conclusion that a residue that moves in counterphase with a group of other residues should not be considered as a part the group, that is, the sign of cross-correlation should not be ignored. As the community analysis for signed networks is still in development (51, 52), we decided to avoid negative links in the protein network by focusing on a rigid body detection approach. For this purpose, we used Local Spatial Pattern (LSP) alignment, a method we developed earlier to discover similar spatial patterns shared by different protein kinases (53, 54). Here, we compared different structures of the same protein to define groups of residues that maintain their mutual orientation during the Molecular Dynamics (MD) simulation. This method represents residues as C_α_-C_β_ vectors and, thus, reflects both translational and rotational degrees of freedom of the residues, providing more strict criteria for protein rigidity. Another advantage of the method is that it uses internal coordinates and does not require global alignment of the protein structures to be compared. As rigid body detection is not limited by residues that are in direct contact, we used extended C_α_ distance cutoff levels ranging from 6Å to 18Å. We suggested that the increase of the cutoff distance between residues can improve sensitivity of the analysis.

The main goal of this study was to find the most informative centrality measures that can detect residues that play important functional or regulatory roles in protein kinases. Calculations were made on non-weighted (NW) networks and with weights calculated by the LSP-alignment and two traditional methods: cross-correlation (CC) and mutual information (MI). Our results show that at short cutoff levels (6Å-8Å) all four methods had rather similar ability to detect important residues in PKA. Increase of the cutoff distance was beneficial for NW and LSP-based networks and detrimental for CC and MI-based networks. In general, LSP-based analysis was outperforming the other three methods. Out of four centrality measures, Closeness centrality was found to be the least effective metrics. Application of Eigenvector centrality was limited due to a small difference between the two largest eigenvalues, also known as spectral gap. We conclude that Betweenness centrality and Degree centrality should be analyzed simultaneously to identify the residues that play important functional or regulatory roles.

## Materials and Methods

### MD simulations

The catalytic subunit of PKA was prepared for all atom molecular dynamic simulations using the crystal structure of wildtype PKA in a ternary complex with Mn2 ATP and inhibitor peptide PKI 6-25 (PDBID: 3FJQ). Mn ions were replaced with Mg and models were processed in Maestro (Schrodinger). Protein Preparation Wizard was used to build missing sidechains and model charge states of ionizable residues at neutral pH. Hydrogens and counter ions were added and the model solvated in a cubic box of TIP4P-EW (55) and 150 mM KCl with a 10Å buffer in AMBER tools (56). Parameters from the Bryce AMBER Parameter Database were used for ATP (57), phosphothreonine (58), and phosphoserine (58). AMBER16 was used for energy minimization, heating, and equilibration steps. Systems were minimized by 1000 steps of hydrogen-only minimization, 2000 steps of solvent minimization, 2000 steps of ligand minimization, 2000 steps of side-chain minimization, and 5000 steps of all-atom minimization. Systems were heated from 0°K to 300°K linearly over 250 ps with 2 fs time-steps and 10.0 kcal*α*mol*α*Å position restraints on protein. Temperature was maintained by the Langevin thermostat. Constant pressure equilibration with a 10 Å non-bonded cut-off with particle mesh Ewald was performed with 300 ps of protein and ligand restraints followed by 300 ps of unrestrained equilibration. Hydrogen mass repartition was implemented to achieve a 4 fs time-step for production runs (59). Production simulations were performed on GPU enabled AMBER16 (60, 61) as above in triplicate for an aggregate of 1.2ms.

### Network creation

#### LSP-based networks

LSP alignment was performed using previously created software (53, 54) adapted for molecular dynamics simulation. Graphs were generated, as described earlier (53), using coordinates of C_α_ and C_β_ atoms for all residues with the exception of glycine (C_α_ and N) and ATP molecule (N_1_,C_8_). In brief: protein structure is represented by a graph with residues as nodes. Links are formed if the distance between C_α_ atoms of the corresponding residues are within a predefined cutoff level ΔC_αα_ (6Å through 18Å in this paper). Each link carries information about mutual orientation of the corresponding C_α_C_β_ vectors: three distances (C_α1_-C_α2_, C_α1_-C_β2_, C_β1_-C_α2_) and the dihedral angle θ (C_β1_-C_α1_-C_α2_-C_β2_). Comparison of two protein structures is presented as a graph with residues as nodes and links created only if all the three distances and the dihedral angles are similar, i.e., within predefined cutoff levels: ΔC_α1α2_<0.2Å, ΔC_α1β2_<0.45Å, Δθ<10°. Weights for the links are calculated using the following formula:
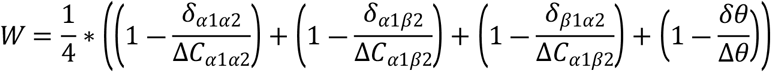
where *δ*_α1α2_, *δ*_α1β2_, *δ*_β1α2_ and *δ*θ are the corresponding differences between two C_α_C_β_ vectors. Weights for the not matching links are assigned zero values.

For LSP-alignment of multiple structures all comparisons were made to a single reference structure and the detected weights were averaged. The reference structure was determined by finding a structure withing the working set that has the smallest C_α_ RMSD to average C_α_ coordinates for the set.

LSP alignment binaries for Linux and MS Windows OS are available free of charge to academic non-profit institutions upon request. For commercial entities please contact UCSD Office of Innovation and Com-mercialization at.

### Non Weighted, Cross-correlation and Mutual information based networks

Cross-correlation and Mutual information matrices were calculated using the Bio3D R package (version 2.4) (62). Three production trajectories 400ns each were aligned by their Ca atoms and merged. Binary contact maps with 7 different cutoff levels (6Å through 18Å) were calculated with the requirement for the residues to be in contact for at least 75% of the trajectory run and used as Non Weighted (NW) matrices. These matrices were subsequently weighted by absolute values of cross-correlation and mutual information.

### Calculation of centralities

Normalized centralities were calculated using igraph R library (version 1.2.5) (63). To calculate betweenness and closeness centralities weights were converted to distances using the following formula: *D*= −log*W*. “Strength” function was used to calculate weighted degree centrality.

## Results

### Scoring key residues

To estimate the efficiency of different metrics in detecting the essential residues of PKA we selected 26 key residues that are highly conserved in the protein kinase family and are known to be involved in different catalytic and regulatory functions (**Figure 1, Table 1**). If a metric would score residues randomly, the accumulated score for 26 residues should be approximately 8% of the total score for the kinase molecule with 336 residues. We, thus, suggested that metrics that can detect important residues should score the selected key residues significantly higher than 8%. **Figure 2** shows the accumulated centrality values for key residues based on four different centrality measures: Degree centrality (DC), Betweenness centrality (BC), Eigenvector centrality (EG), and Closeness centrality (CL) calculated with seven different distance cutoff values. Several observations can be made from these data. Closeness centrality proved to be the least effective metrics. CL scores were almost identical in all cases with the key residues share between 8% and 9%, indicating that Closeness centrality consistently failed to distinguish the key residues from the rest of the molecule. The most effective metrics was Betweenness centrality in all four types of the networks with its efficiency rising with increasing cutoff levels in Non-Weighted and LSP-based networks. Eigenvector centrality revealed a non-consistent pattern, especially in the LSP-based networks changing from nearly zero values to about 12%. Eigenvector centrality, by definition, is a measure of the influence of each node. Nodes have a high influence if they are connected to other influential nodes. At the same time, the influence of the neighbors also depends on the influence of the original node. This recursive problem is solved mathematically by finding eigenvectors of the adjacency matrix and associated with them, eigenvalues, hence, the name of the centrality measure. The problem is that eigenvector centrality is always associated with the first/largest eigenvalue, which is suggested to be much higher than the other eigenvalues. For example, in the case of Imidazole glycerol phosphate synthase, the authors reported that the first eigenvalue was two orders of magnitude higher than the rest of the eigenvalues (28). Our analysis of eigenvalues in all the networks (**Figure S1**) showed that in all cases first two eigenvalues were very close especially in the CC and MI-derived networks and smaller cutoff levels for all networks. This is an apparent result of the bilobal shape of the kinase molecule that led us to the conclusion that in our case Eigenvector centrality is not a suitable method.

**Table 1.**
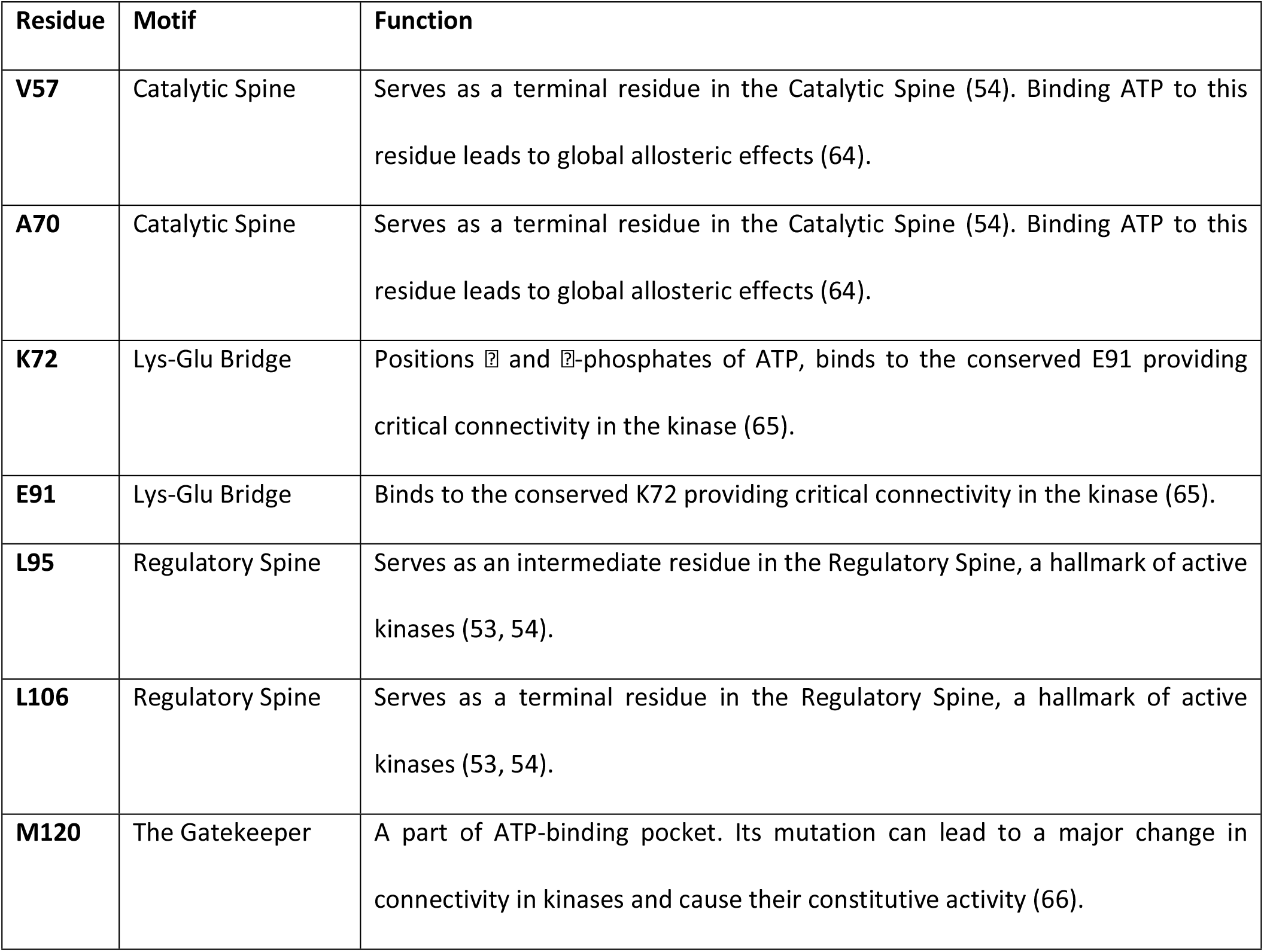
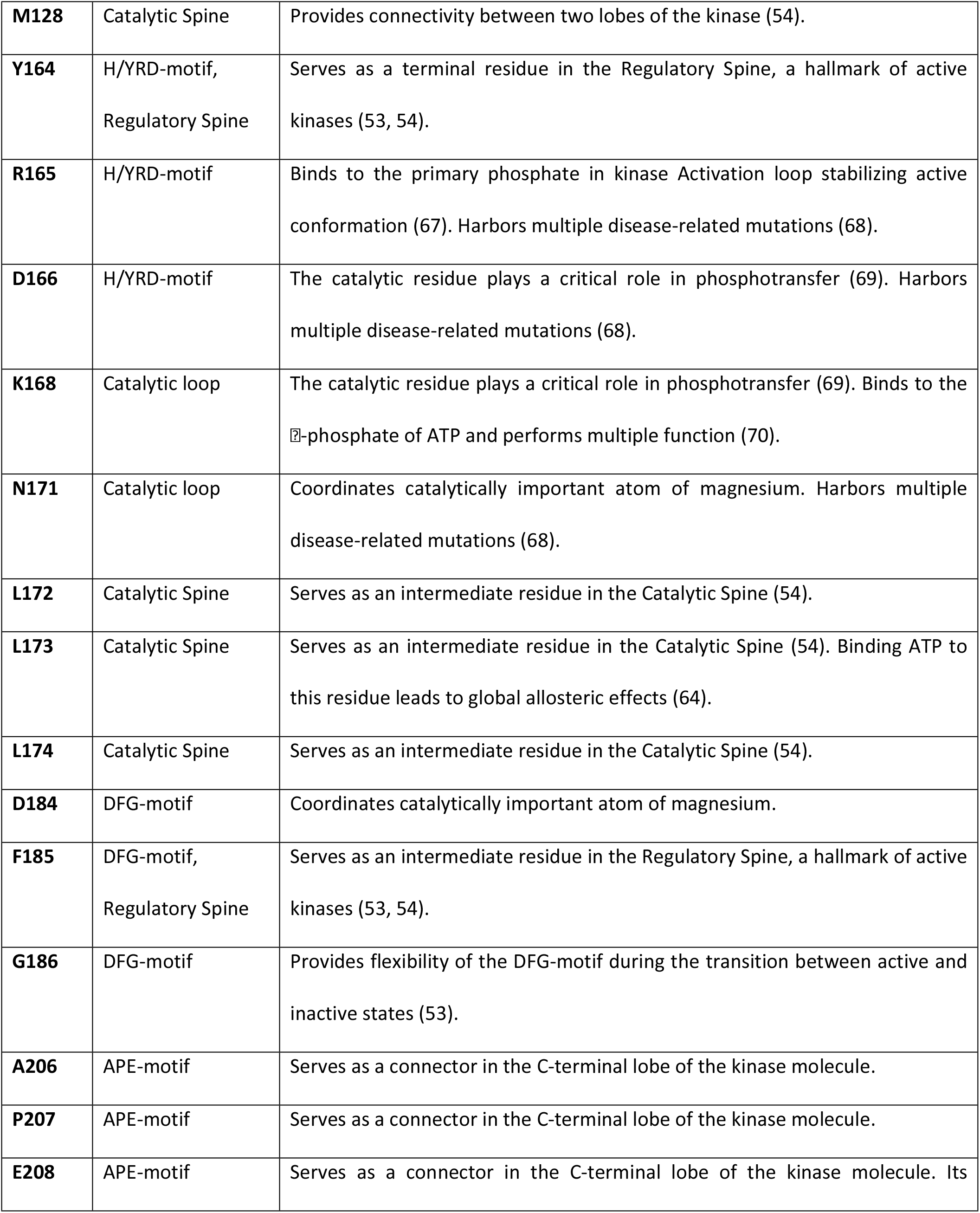
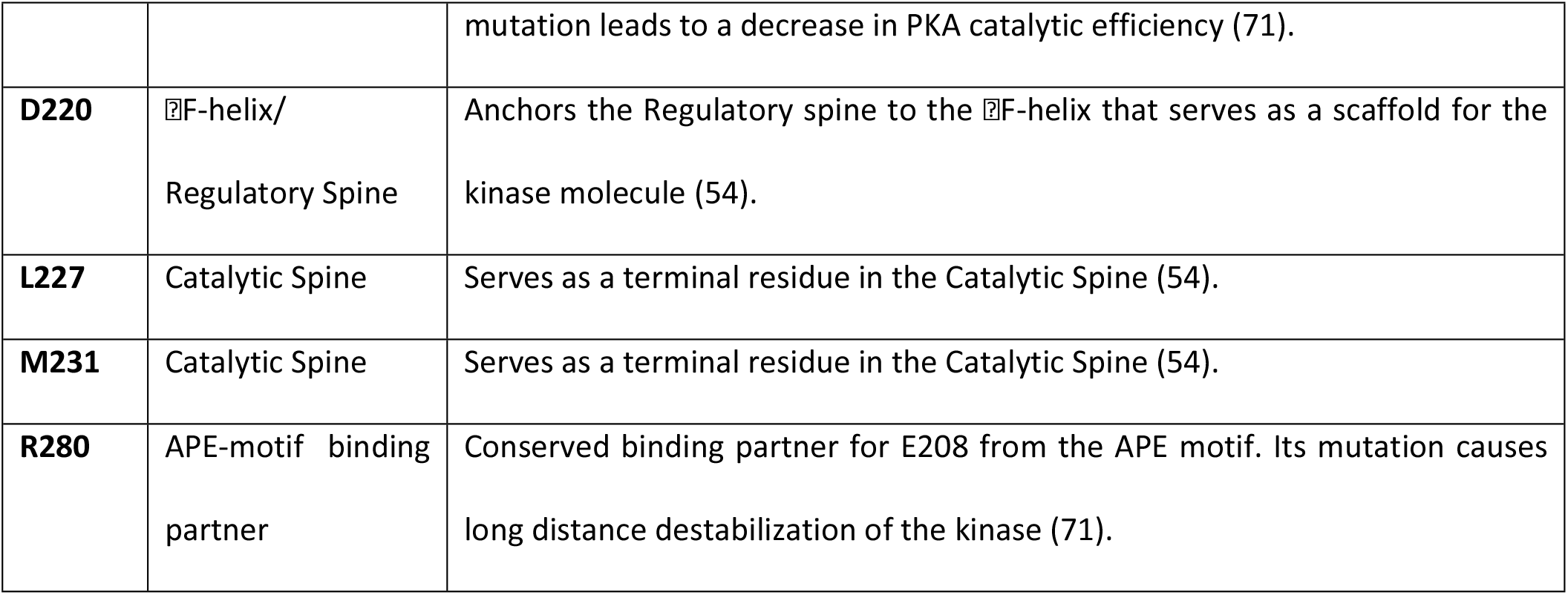
Description of the 26 key residues selected for centrality scoring presented in Figure 1.

**Figure 1.**
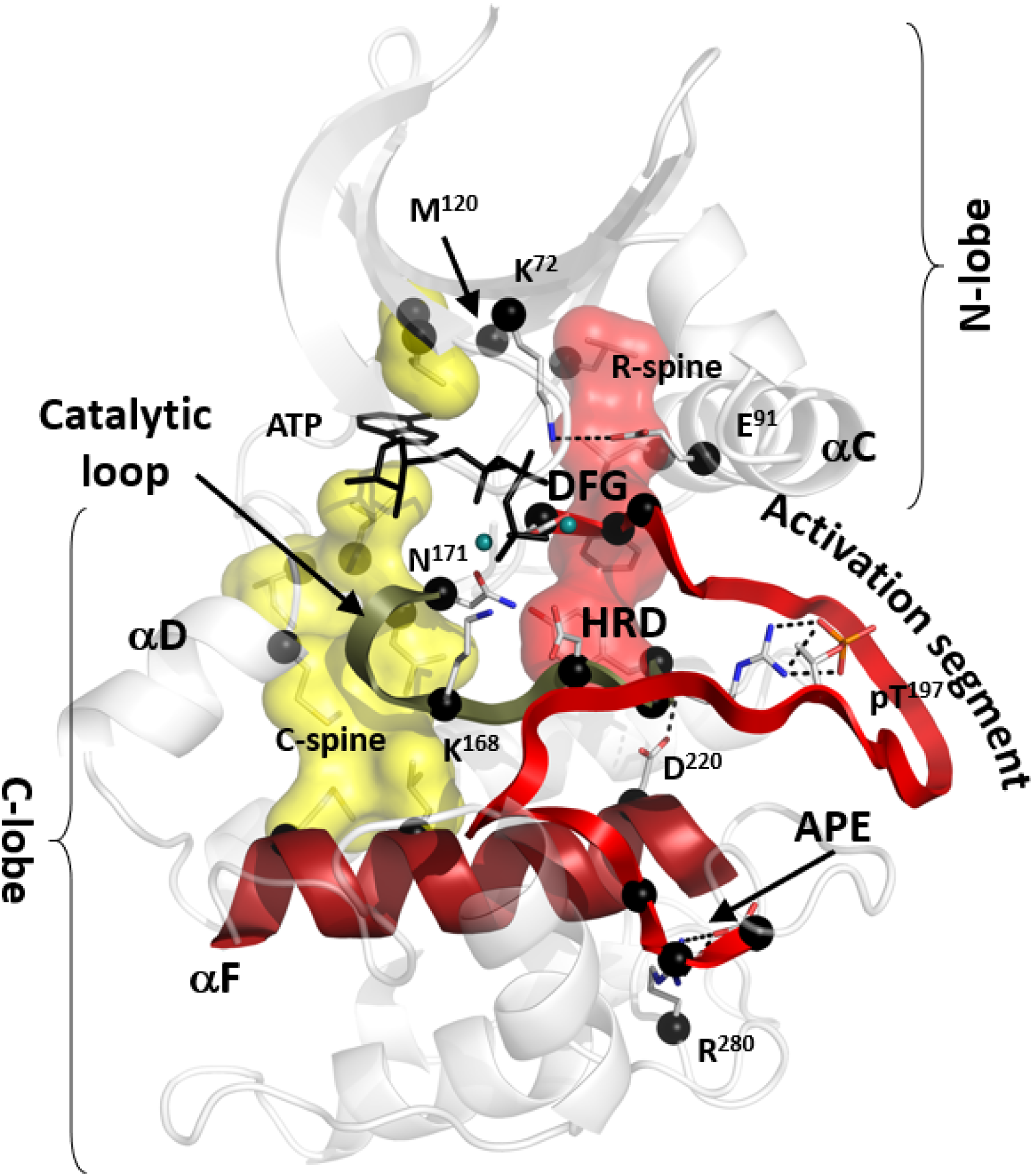
Conserved core of the eukaryotic protein kinases with the 26 preselected key residues (black spheres). The core consists of two lobes: N-lobe and C-lobe. ATP (shown as black sticks) is sandwiched between the lobes bound to two magnesium ions (teal spheres). Two hydrophobic ensembles C-spine (yellow surface) and R-spine (red surface) span the core providing global connectivity within the molecule. DFG-motif and APE-motif are flanking the Activation segment (red ribbon). HRD-motif is a part of the Catalytic loop (olive ribbon). Both spines are anchored to the ⍰F-helix (dark red) that spans the C-lobe and serves as a foundation for the catalytic machinery of the kinase. Additional description of the selected key residues is provided in the **Table 1**.

**Figure 2.**
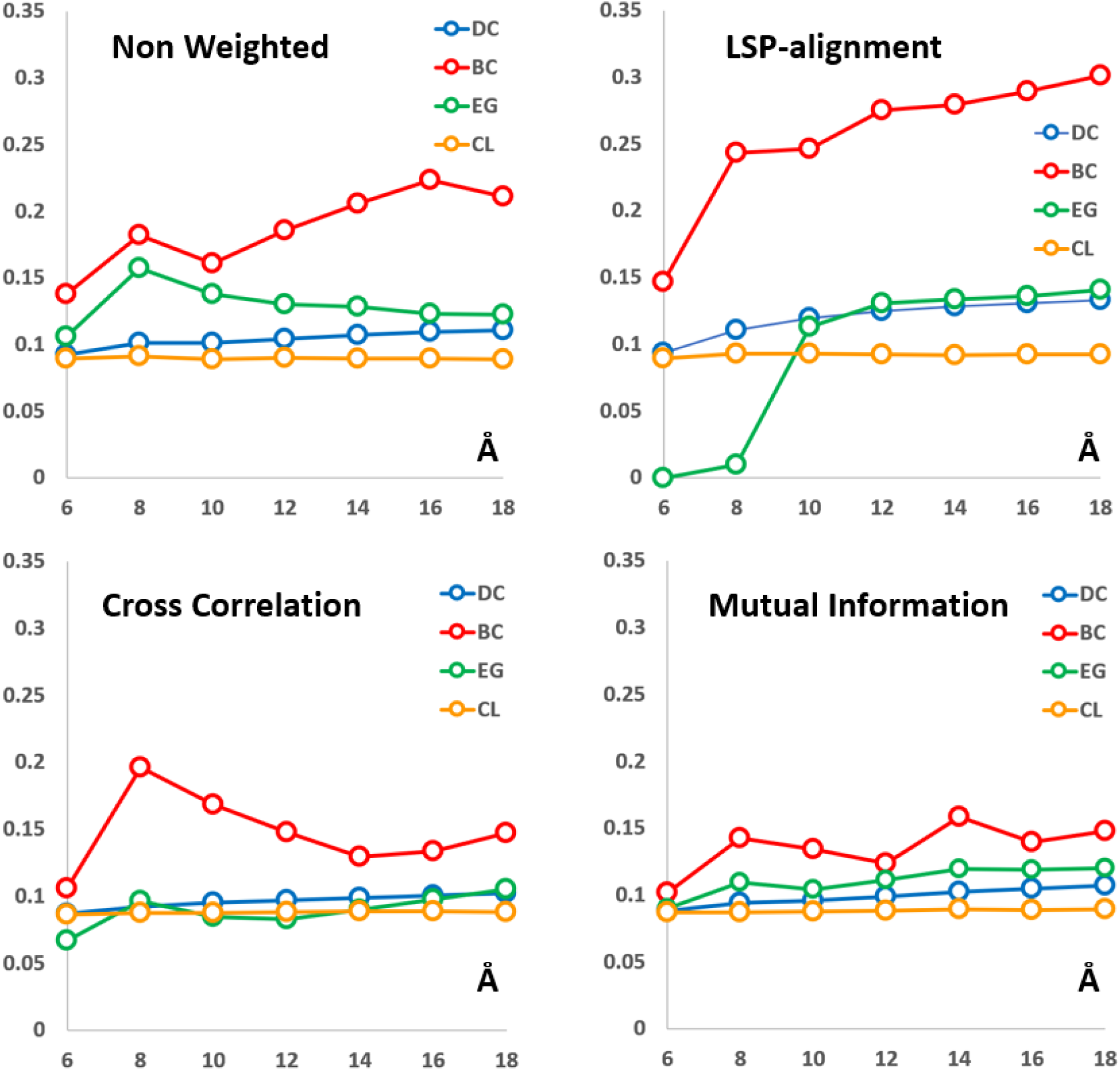
Accumulated scores for the 26 key residues defined by four different metrics: Degree centrality (DC), Betweenness centrality (BC), Eigenvector centrality (EG) and Closeness centrality (CL) using different ΔC_αα_ cutoff levels (X-axis). Four different methods were used for network constructions: binary (Non Weighted) networks, Local Spatial Pattern alignment, Cross correlation, and Mutual Information. See Materials and Methods for details.

### Role of the trajectory length

Proteins in general, and protein kinases in particular, are known to be highly dynamic objects with timescales for different motions from femtoseconds to seconds. How stable the detected residue networks are is an open question and weather our results can benefit from longer simulations, or they can be derived from shorter trajectories? To evaluate the reliability of our results and its dependence on the trajectory length we calculated Degree and Betweenness centralities for all PKA residues using trajectory fragments ranging from 1ns to 120ns. Calculations were repeated ten times using neighboring fragments of the trajectory and the replications were compared to each other. Average correlation coefficients between the replications for each interval are shown in **Figure 3**. Non-Weighted networks showed a very high level of consistency with small standard deviation values, although a noticeable decrease can be observed after 20ns. Introduction of weights significantly reduces the reliability especially for the Betweenness centrality calculations. This is an expected result as Betweenness centrality is a global parameter and a single change of connectivity between two nodes can lead to changes of betweenness throughout the whole network. Degree centrality, on the other hand, is a local parameter and changes of connectivity have no global effect. 1 ns long trajectories produced the least reliable results with reliability gradually increasing up to 10-20 ns. After that the reliability either plateaued or decreased with higher levels of standard deviation. This allowed us to conclude that trajectory length around 10-20 ns is the optimal duration for the current analysis.

**Figure 3.**
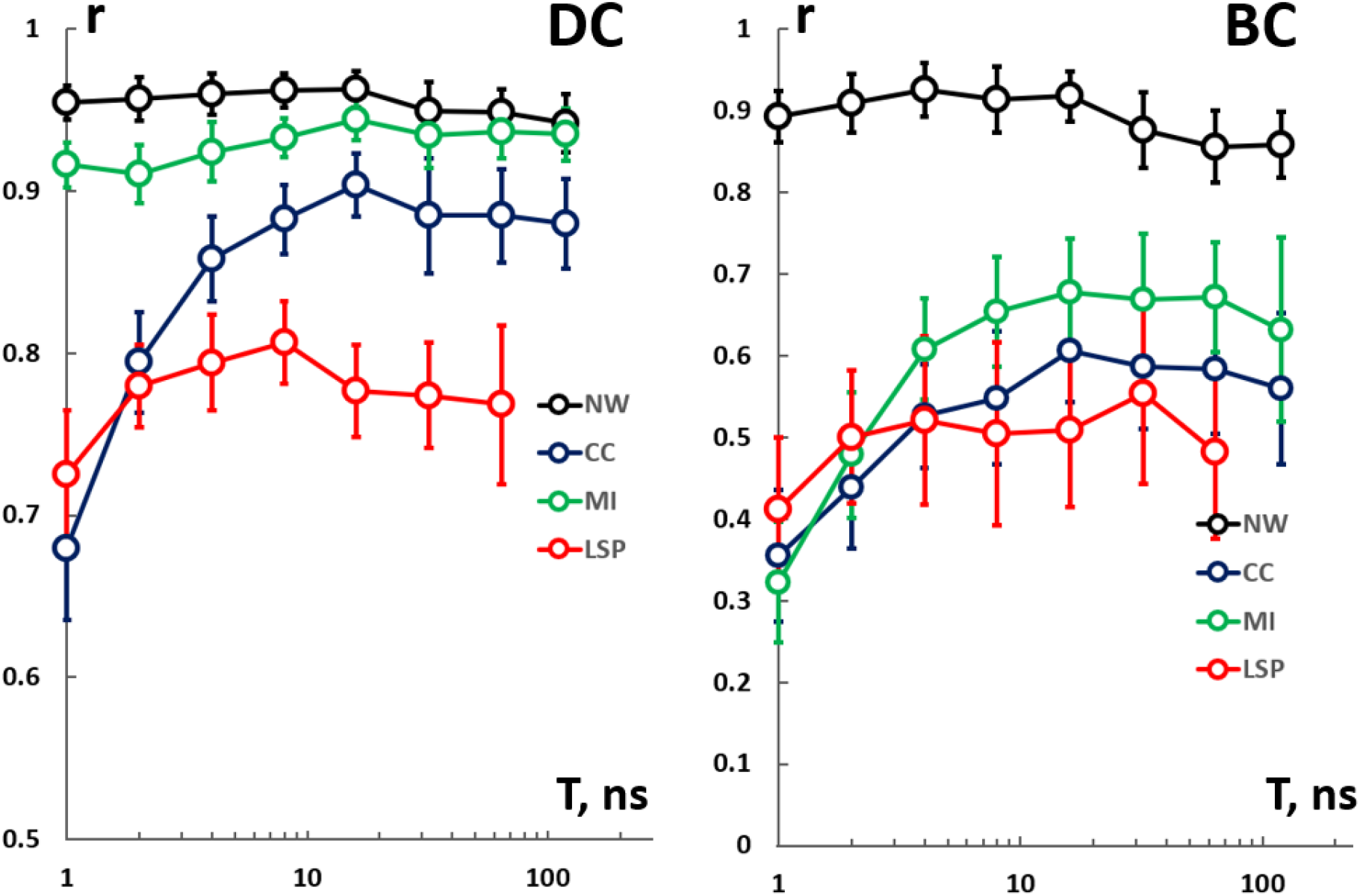
Average correlation coefficients (r) between 10 replications of centrality measures calculated for different lengths of MD trajectories (T). Degree centrality (DC) and Betweenness centrality (BC) were determined for Non-weighted (NW) networks, Cross-correlation (CC), Mutual information (MI) and LSP-based networks. Standard deviations for each set of 10 replications are shown as error bars. (See text for details).

### Visualizing centralities in PKA

We used ten 16ns long intervals to calculate four centrality measures for the whole PKA molecule using 8Å cutoff level for CC and MI-based networks and 12Å cutoff level for NW and LSP-based networks and mapped the average values on the PKA structure (**Figure 4**). Again, as the results for the key residues indicated (**Figure 2**), Closeness centrality was the least informative parameter as the distribution of its values was virtually identical in all four networks with high level of CL concentrated in the middle of the molecule. Eigenvector centrality values distribution also confirmed our suggestion that its use is hindered by the bilobal architecture of the kinase molecule: all highly scored residues in NW and LSP-based networks were located in the C-lobe, while in CC and MI-based networks they were in the N-lobe. As it was mentioned earlier, Degree and Betweenness centralities characterize distinctively different network properties. While DC reflects local connectivity, BC identifies critical communication points between densely interconnected areas. As DC and BC reflect two different topological characteristics of networks, we visualized them simultaneously as a scatterplot (**Figure 5**).

**Figure 4.**
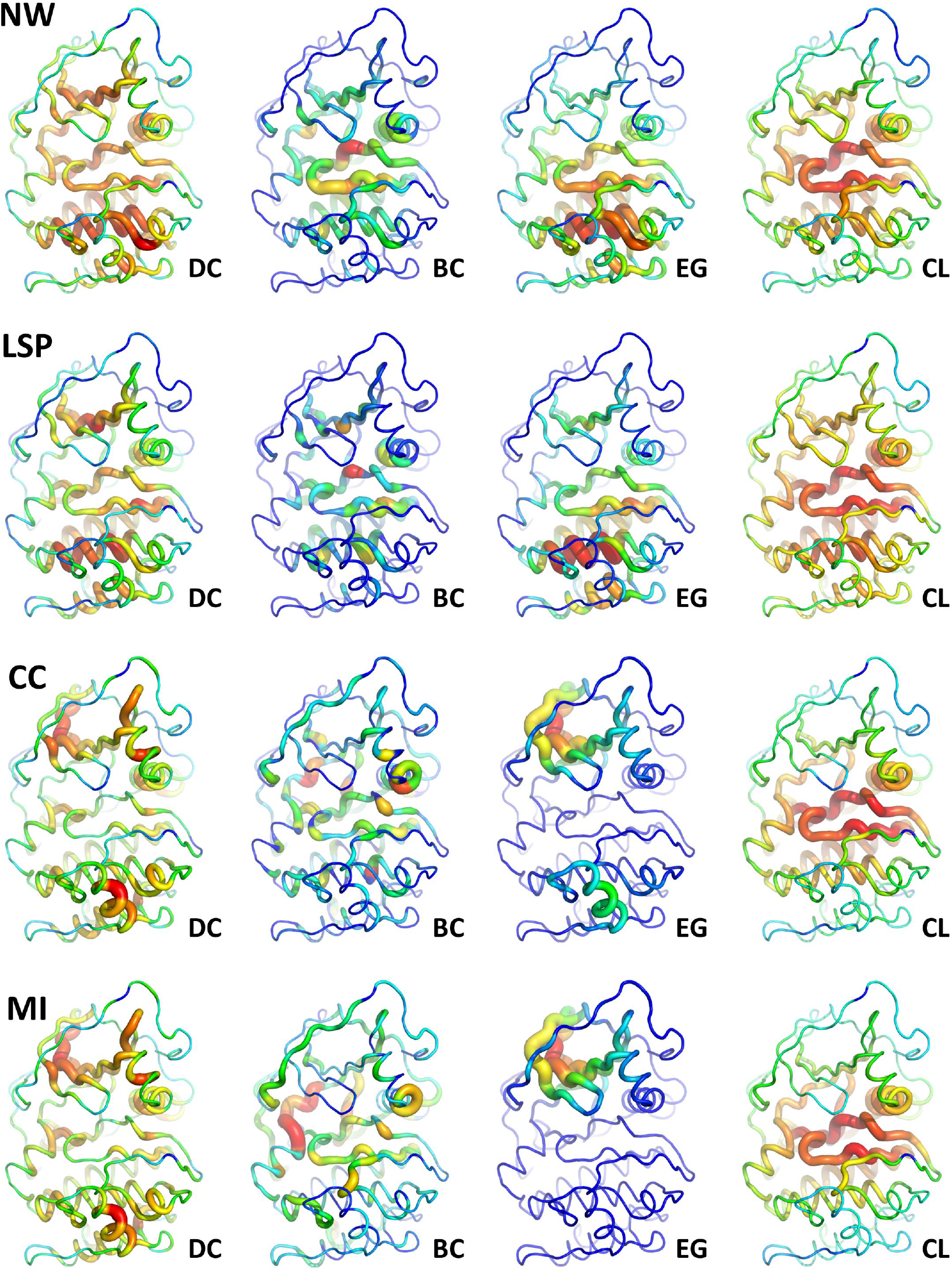
**Four centrality measures Degree centrality (DC), Betweenness centrality (BC), Eigenvector centrality (EG) and Closeness centrality (CL) calculated in four residue networks: Non-weighted (NW) and with weights calculated using LSP-alignment (LSP), Cross-correlation (CC) and Mutual information (MI).** Centrality values are mapped on PKA structure (1ATP) using Pymol visualization program. Calculations were made using 12Å DC_aa_ cutoff level for NW and LSP and 8Å for CC and MI networks. Centralities were calculated for ten neighboring 16ns intervals and averaged.

**Figure 5.**
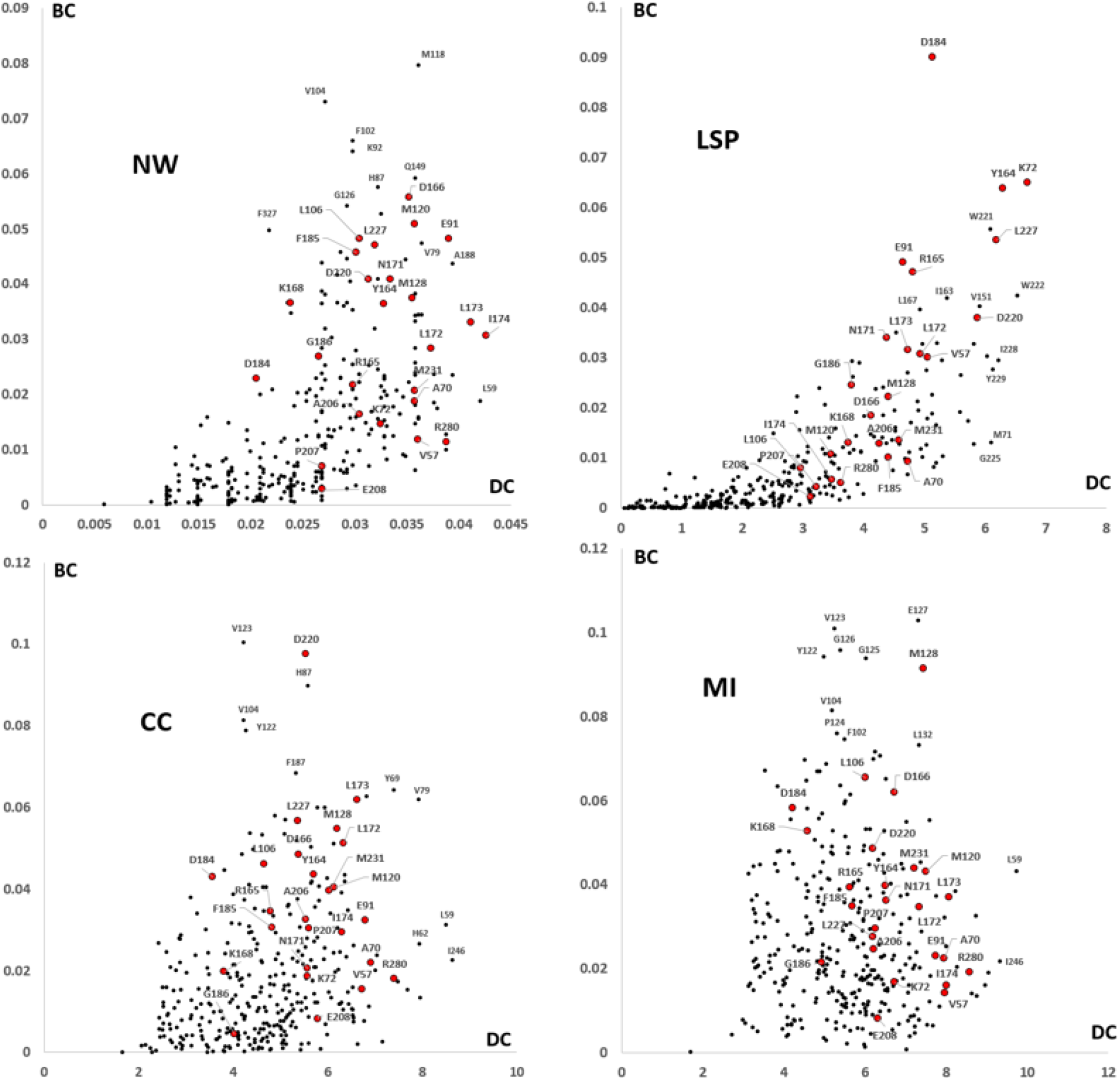
Scatterplots of Degree centrality (DC) and Betweenness centrality (BC) in PKA residue networks: Non-weighted (NW) and with weights calculated using LSP-alignment (LSP), Cross-correlation (CC) and Mutual information (MI). Red dots represent 26 key residues shown on **Figure 1**. Calculations were made as indicated in the **Figure 4** legend.

In general, distribution of the 26 key residues on the plots with respect to the rest of PKA residues confirmed our preliminary estimate efficiency of the four methods to detect the most important residues (**Figure 2**). LSP-based provided a clear distinction of 6 well-recognized residues including the D184, responsible for phosphotransfer, universally conserved Lys-Glu bridge (K72-E91) that stabilizes active conformation in protein kinases, two residues from the H/YRD motif (Y164 and R165) and L227 that connects the C-spine to the F-helix. Other highly scored residues included a conserved W222, that connects the APE-motif to the F-helix (See Fig.5 in ref. (54)). Noticeably, the APE-motif itself received an average level of DC and very low BC. Non-weighted network also scored E91 and L227 rather high, however, very high scores were assigned to residues that are not known to be involved in any critically important functions. CC and MI-based networks produced even less informative plots with the known important residues scattered in the middle of the widely distributed points.

## Discussion

Complex network studies belong to a young and quickly developing field of science with a long list of questions still waiting to be answered. Statistical significance of network characteristics (72, 73) and properties of networks with negative connections (74, 75) are, to name a few, among most essential problems. Consequently, the results presented here are far from the definitive solution of the network centrality problem in proteins. Nevertheless, they allowed us to cast light on important issues of the problem, providing a foundation for future development. Most importantly, we questioned the process of protein network creation. Currently links in these networks are based on the concept of interaction between the residues, i.e., a link is created if two residues are in direct contact. We argued that the links should not represent simple interactions but should reflect conservation of distance between two residues in time. In other words, tightly interconnected parts of the residue network should correspond to dynamically stable, semi-rigid regions, similar to tight social communities that share a particular characteristic/interest/activity. This approach is consistent with community analysis of protein dynamics (40) where often communities are viewed as semi-rigid bodies that persist during the simulation (48-50). This conceptual change led us to the following assumptions: first – the cutoff level for distance between two linked residues does not need to be restricted by direct contacts and can be increased to the size of the community. Second – negative and positive correlations for residue motions cannot be merged to define link weights as it is done in two popular methods: Cross-correlation and Mutual Information (40). We argued that if two residues move in an anti-correlated way, they should belong to different communities, similar to the social networks with “friends” and “foes”, i.e., negative links (74, 75).

To define link weights that are consistent with our hypothesis, we used the LSP-alignment, a method that can detect similar spatial patterns formed by C_a_C_b_ vectors in two different protein structures. We compared these networks to binary (non-weighted) networks derived from the same trajectories and to networks with weights defined by the two traditional methods: Cross-correlation and Mutual Information. We then compared four popular centrality measures to evaluate their efficiency in finding important residues in PKA.

In this work we were facing a complicated “bootstrapping” problem: to find the most efficient method that can detect important residues without preliminary knowledge about either the method efficiency or the residues to be detected. To resolve this problem, we compiled a set of 26 residues in PKA that are known to be involved in different catalytic and regulatory functions (**Figure 1, Table 1**). There was no guarantee that all of these residues would have high centrality values, or the set provides a comprehensive coverage of all important residues. Nevertheless, we hypothesized that there is a high probability that a set of important residues that can be detected by graph theoretical methods and the set of 26 known key residues can overlap. We, thus, expected that, in average, an efficient method should be able to distinguish the preselected set from the rest of the molecule.

In general, the results presented on **Figure 2** were consistent with our hypothesis: increase of the cutoff level was beneficial for non-weighted and LSP-based networks and detrimental for CC-based networks or not effective for MI-based networks. It is rather expected result as larger cutoff level should introduce more negative correlations into the networks. The CC and MI-based networks, as expected had the best performance at the well-known 8Å cutoff level that roughly corresponds to the direct contacts between residues.

Out of the four most popular centrality measures we excluded Closeness centrality that was not capable of distinguishing known important residues from the rest of the molecule (**Figure 2**) and revealed almost identical distributions in the whole PKA molecule (**Figure 4**). The remaining three centralities are often discussed in the literature together. Degree centrality (DC) and Betweenness centrality (BC) are the most basic and the oldest measures that have opposite natures: while DC is a local measure of connectivity, BC is a global characteristic that reflects a flow of information inside the network. Eigenvector centrality (EG) was conceived as a compromise between these two parameters as it reflects both local and global properties of the network (76). Several reports indicated that the EG can be an efficient tool that can find functionally important residues (28, 77). Nevertheless, we could not use EG in the case of PKA due to the narrow spectral gap of the network (**Figure S1**). Spectral gap is a difference between the two largest eigenvalues that is required to be large for EG calculation. Physically the small spectral gap means that the network has a clearly defined “bottleneck” separating the network into relatively isolated parts (78). In the case of protein kinases, the observed small spectral gap is an obvious consequence of their bilobal structure that makes the application of EG for protein kinases questionable in general. The same problem can occur for any proteins or protein complexes that do not have an overall globular architecture.

Although Betweenness centrality, in the LSP-based networks in particular, was very effective in discriminating the key residues from the rest of the molecule (**Figures 2**,**4**), it had the highest level of noise (**Figure 3**) that can be attributed to the global nature of this metrics. Another factor is dynamic changes in the residue networks that currently are not well understood and will require additional studies. Our results show that while increase of simulation time from 1ns to around 20 ns is beneficial, the following increase up to a microsecond timescale did not provide any improvement of the network centralities calculation. A possible alternative to Betweenness centrality can be Flow Betweenness (9) that can be less sensitive to local changes in connectivity during the simulation.

Despite the presence of the significant noise in the BC values in the LSP-based networks, the method could rather efficiently separate several important PKA residues from the rest of the molecule (**Figure 5**). Importantly, the nodes can be roughly separated into two groups of residues, one group has a high level of DC and a low level of BC. These residues are the most interconnected nodes that do not participate in global information transfer. They are in the center of tightly bound regions either in the N-lobe (like M71) or the C-lobe (G225, I228, Y229). The second group has a high level of DC and also a high level of BC. These resides can be viewed as super communicators in the network. Not surprisingly residues from this group are well-known members of the major protein kinase motifs: HRD, DFG, Lys-Glu bridge, and Catalytic spine.

These results are in line with similar studies of protein interaction networks in yeast where concepts of “date-hubs” and “party-hubs” were introduced (79). The former are nodes with high BC and high DC levels, while the latter are nodes with low BC but high DC values. Such distinction proved to be useful in the characterization of proteins essentiality and expression dynamics (4, 80). This is another example of the universal nature of topological properties that can pinpoint important parts of a network irrespective of its physical nature. However, as our results indicate, the network creation step can be crucially important for the efficiency of this approach.

Admittedly, these results were obtained using a single protein, albeit representing and important protein family. Several essential questions also remain to be answered. Does Eigenvector centrality in proteins with a large spectral gap provide any advantage in comparison to Degree or Betweenness centralities? What is the nature of slow sub-microseconds timescale drifts that change connectivity in residue networks? Nevertheless, the observed strong performance of the LSP-derived networks is encouraging and provides foundation for the further development of the graph-theoretical analysis of protein networks.

## Conclusions

Based on the idea that residue networks have to reflect stable, semi-rigid regions in the protein, we argued that currently accepted practice of using cross-correlation and mutual information for defining link weights should be reconsidered. We showed that the corresponding change of the network construction method was beneficial for detection of functionally important residues in protein kinase A, which was used as a well-studied benchmark. We recommend calculating two centrality measures: Betweenness centrality and Degree centrality and plot the amino acid residues on a joint scatterplot. Residues with high level of BC and high level of DC are predicted to be major communicators within the protein structure. Mutations of these residues should lead to global changes of protein dynamic landscape. Residues with high level of DC but low level of BC are predicted to be local centers of densely interconnected communities with low conformational entropy. These predictions can be tested experimentally via biochemical and NMR studies.

## Author Contribution

APK designed the research, performed calculations, and wrote the manuscript. PCA performed MD simulations. APK, PCA and SST collectively evaluated the ongoing calculations and edited the manuscript, SST provided financial support for the project.

## Acknowledgements

The authors thank Dr. Michal Vieth (Lilly Biotechnology Center) for fruitful discussions during preparation of the manuscript. The authors thank Evan K. Kobori (UCSD) and Dr. Ruben Abagyan (UCSD) for their thoughtful feedback on the manuscript. The work was supported by Eli Lilly Research Award program and the NIGMS, National Institute of Health (Grant 5R35GM130389 to SST). The authors declare no conflicts of interest.

## Supplemental information

**Figure S1.**
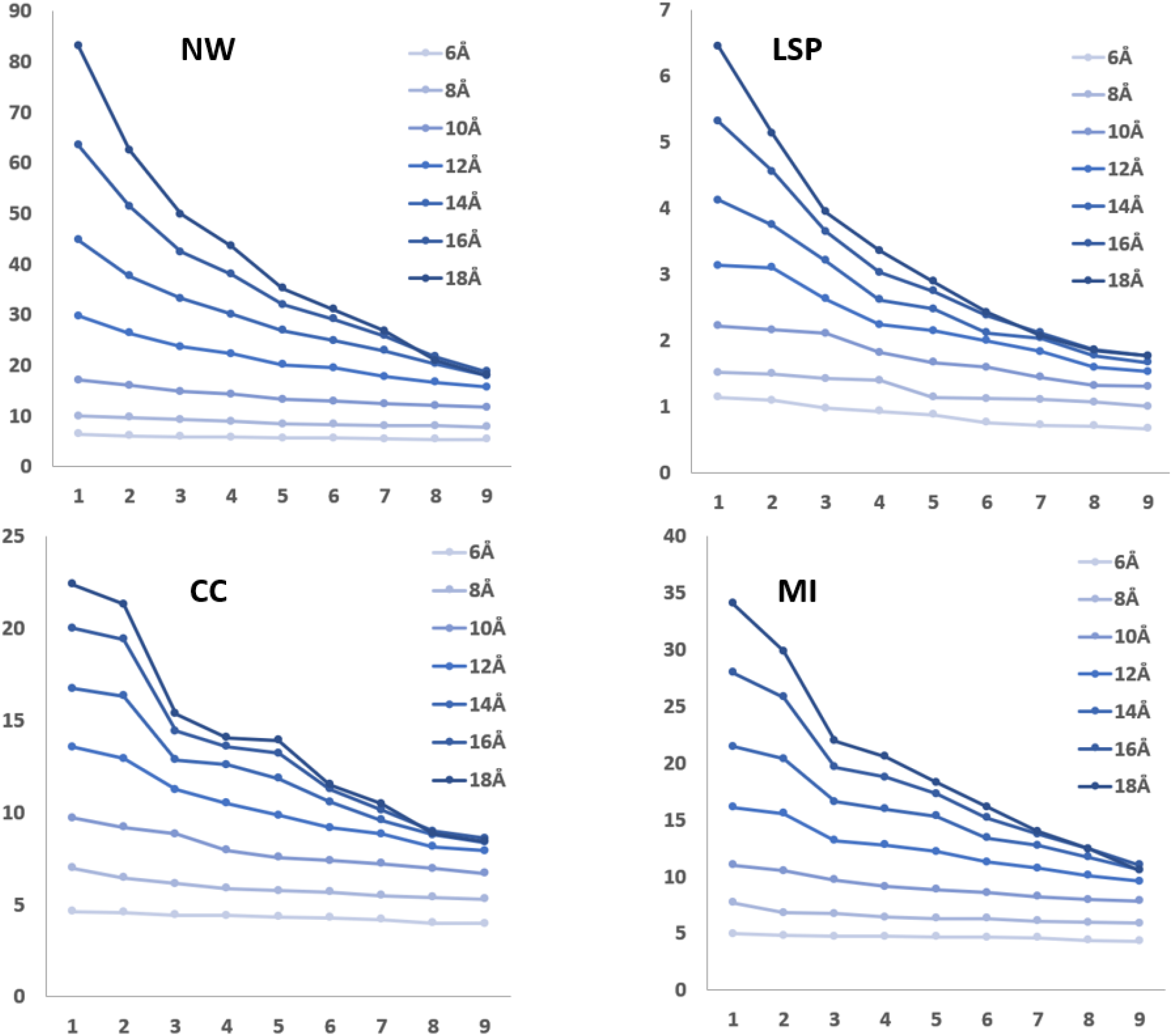
Nine largest eigenvalues for different residue networks created using seven level DC_aa_ cutoff levels. Calculations were made for total 1.2ms trajectory.

